# Anticipatory Control of Momentum for Bipedal Walking on Uneven Terrain

**DOI:** 10.1101/769828

**Authors:** Osman Darici, Hakan Temeltas, Arthur D. Kuo

## Abstract

Humans and other walking bipeds often encounter and compensate for uneven terrain. They might, for example, regulate the body’s momentum when stepping on stones to cross a stream. We examined what to do and how far to look, as a simple optimal control problem, where forward momentum is controlled to compensate for a step change in terrain height, and steady gait regained with no loss of time relative to nominal walking. We modeled planar, human-like walking with pendulum-like legs, and found the most economical control to be quite stereotypical. It starts by gaining momentum several footfalls ahead of an upward step, in anticipation of the momentum lost atop that step, and then ends with another speed-up to regain momentum thereafter. A similar pattern can be scaled to a variety of conditions, including both upward or downward steps, yet allow for considerably reduced overall energy and peak power demands, compared to compensation without anticipation. We define a “persistence time” metric from the transient decay response after a disturbance, to describe how momentum is retained between steps, and how far ahead a disturbance should be planned for. Anticipatory control of momentum can help to economically negotiate uneven terrain.

## Introduction

Humans and other bipeds often encounter uneven terrain in walking environments. Even the act of stepping between curb and sidewalk can perturb otherwise steady walking and cause a loss of stability and forward speed. Although gait may be restored with feedback control, there may be advantages to anticipating the upcoming disturbance, for example through vision or other imaging, and performing compensations before stepping on it. Although it may seem advantageous to anticipate ahead of time, it is unknown how far ahead to look, what actions to take, and at what cost to restore nominal gait. A simple modeling analysis can yield insight on how best to anticipate and compensate for a terrain disturbance.

Anticipation offers a potential advantage in time. Whereas a feedback response cannot be initiated until a disturbance has been physically encountered, anticipatory control can be initiated immediately upon its (visual) detection. It can thus distribute control actions both before and after the disturbance, and potentially compensate more economically or with less actuator effort. A simple example is the running long jump, which builds up more momentum and yields a longer jump than can be gained from a static, standing position. Similar benefits might apply to traversal of a minor terrain disturbance, with appropriate anticipation and planning.

We will address anticipation and integration as a simple optimization problem (Fig. 1). One simplification is to consider only a single disturbance in terrain height (“up-step” or “down-step”). Many a person has experienced a stumble or fall from a small, unexpected disturbance, as has many a bipedal robot. In fact, an up- or down-step is often used as a test of a robot’s robustness to uneven terrain^1,2^. An additional simplification is to consider a minimal, sagittal plane model of bipedal walking, in which momentum is carried by a pendulum-like stance leg, and additional degrees of freedom for the swing leg or lateral motions are ignored (or controlled by lower-level feedback loops). Pendulum-like, underactuated locomotion describes some aspects of human walking e.g.^3,4^, and that of a number of robot models^5–7^ and physical machines^8–12^. Underactuation implies that energy change occurs little during stance, but substantially with a relatively rigid swing leg’s collision with ground^13–15^, as part of the “step-to-step transition”^16^. The energy and momentum losses from collision, and the positive work done to offset these losses, are key to the optimization solutions.

**Figure 1.**
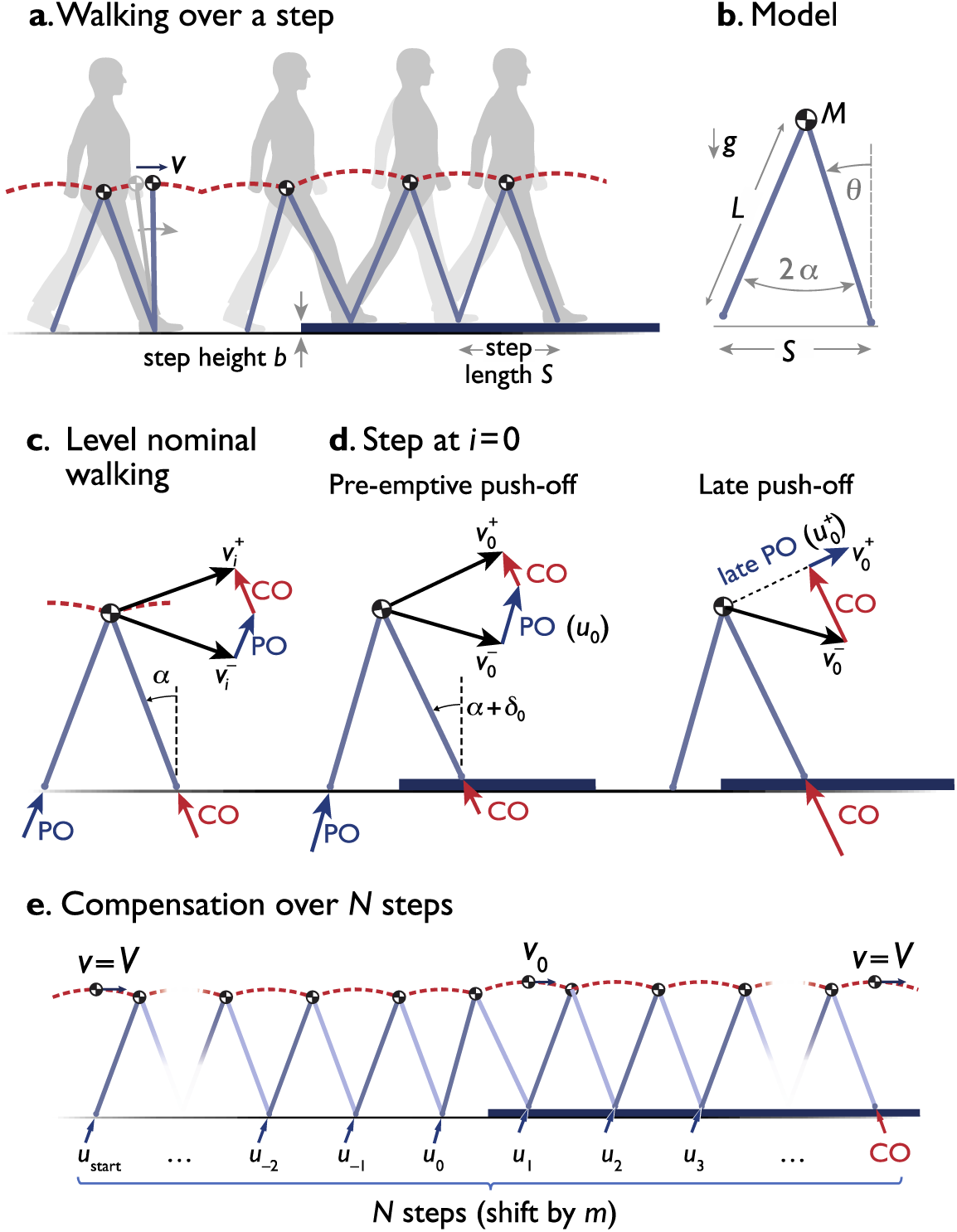
Simple model of walking over a single up- (or down-) step. (**a**) Body center of mass (COM) is a point mass carried atop stance leg acting as an inverted pendulum, encountering step of height *b*. (**b**) Model has mass *M* at pelvis, massless legs of length *L*, gravitational acceleration *g*, and fixed step length *S*, with inter-leg angle 2*α*. Stance leg angle is measured with angle *θ* as shown. (**c**) Level nominal walking has step-to-step transition where COM velocity is redirected from forward-and-downward 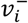 to forward-and-upward 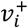, with active, impulsive trailing leg push-off (PO), followed by inelastic, impulsive, leading leg collision (CO). (**d**) Up-step occurs on step *i* = 0, with two cases: pre-emptive and late push-off, which both perform positive work (*u*_0_ and 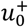, respectively) but earlier or later than collision. (**e**) Optimization is to modulate *N* push-offs to minimize their total work while accommodating the up-step in the same time as level walking at speed *V*. The *N* steps are nominally centered on the up-step for equal anticipation and recovery, but may be shifted by *m* steps relative to obstacle. Up-step heights have positive *b*; down-steps negative *b*.

The proposed optimization modulates forward momentum to mitigate the effects of a single terrain height disturbance (an “up-step” or “down-step”). The goals will be to regain a nominal walking gait without falling down, and without loss of time, so that the biped will eventually be at an appointed time and position, as if had always been on level terrain. Of course, there may be some demands on overall or peak work of muscles or actuators, and so these will be minimized or evaluated as part of the optimization. Here we present the dynamics of a simple model (Fig. 1), followed by an optimization formulation to address these questions. This is then followed by a series of simple observations regarding how to anticipate and recover from a terrain disturbance.

## Results

We found substantial advantages to anticipating and compensating for an up-step. We first found that without compensation, an up-step causes a considerable loss of time. The lost time may then be regained by speeding up from nominal walking, but even the optimal speed-up places considerable demands on overall work and peak actuator power. We then found that the optimal compensation is instead to anticipate and adjust forward momentum or speed in a transient period starting before and ending after the up-step. Results show that a single compensation pattern can be scaled to characterize optimal strategies for different walking speeds and step heights, including down-steps.

### Uncompensated transient response resembles exponential decay and is described by persistence time

The benchmark for optimal compensation strategies is the effect of a completely uncompensated up-step. Uncompensated control means that only the nominal, pre-emptive push-off is applied^17^ (*u*_*i*_ = *U*), without any modulation. The transient response resembles an exponential decay, starting with an immediate loss of speed at the up-step, followed by an approximately exponential response over subsequent discrete steps (step index *i*; equation 2). We defined the time constant of the exponential decay as a *persistence time* (see Methods), to characterize how compensations scale over a wide range of walking speeds and disturbances.

There is considerable advantage to pushing off onto the up-step pre-emptively. If the up-step can be anticipated only enough to allow push-off to occur before the collision, it can greatly reduce the immediate loss of speed compared to a late push-off^17^. For a sample up-step of modest height (*b* = 0.025*L*), loss of speed is about 80% less and time loss is 82.7% less (see Fig. 2a-b). Continuing with nominal push-offs after the up-step, both cases experience a similar type of exponential decay in speed error, and therefore an asymptotic return to nominal walking (Fig. 2b). But pre-emptive push-off reduces the total time loss by 82% compared to late push-off (for *b* =0.025*L*), even if push-off magnitude is not actively modulated. Thus, even a short-range ability to detect an upcoming obstacle can be helpful, if it allows for an unmodulated but still pre-emptive push-off.

**Figure 2.**
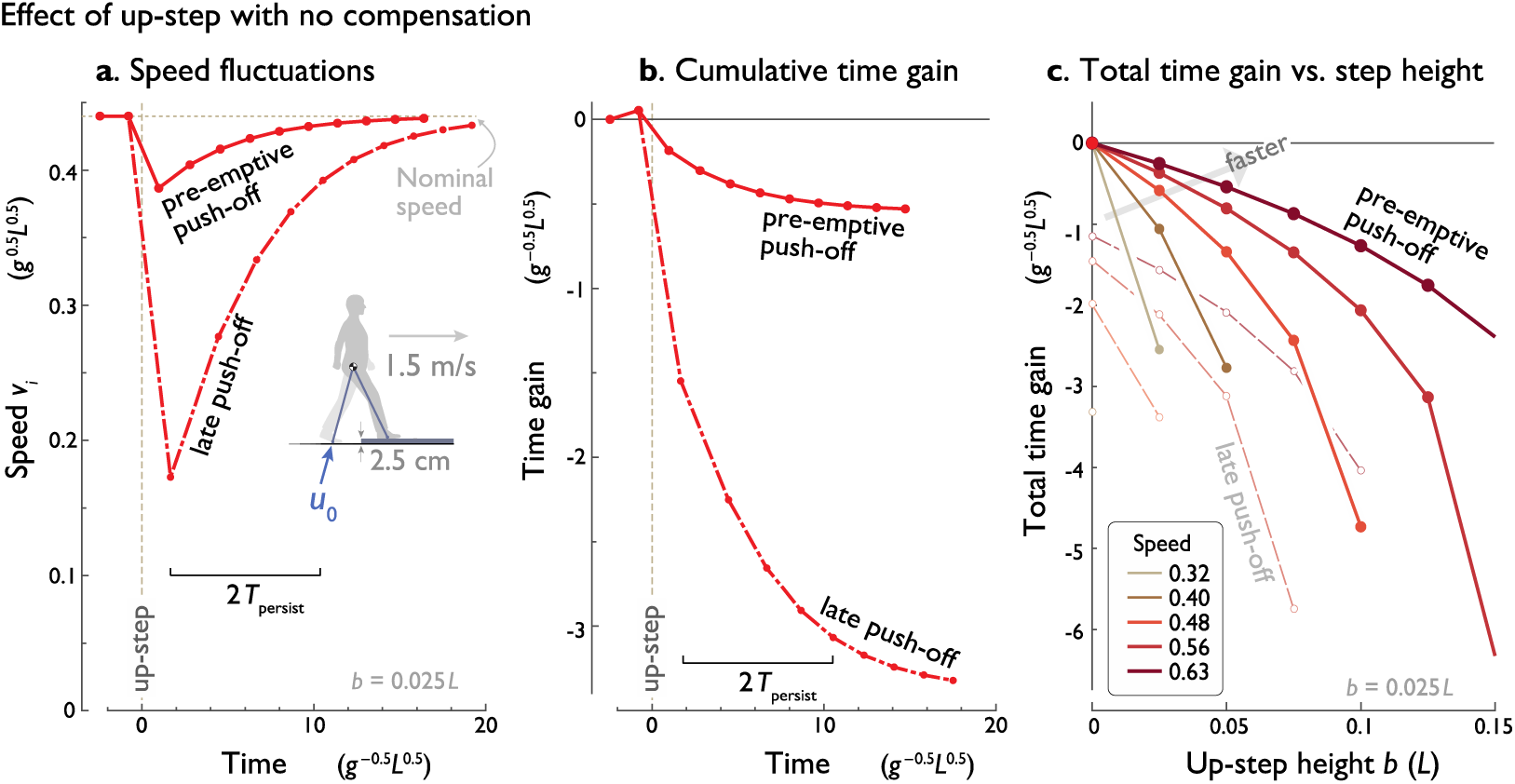
Effect of an uncompensated up-step. (**a**) Speed fluctuations vs. time, for an up-step at time 0 with constant push-off. Model experiences transient disturbance from nominal speed due to up-step, with smaller effect if push-off occurs pre-emptively as opposed to late. Each dot (filled symbol) indicates a discrete step, with speed *v*_*i*_ sampled at mid-stance (see Fig. 1e). (**b**) Cumulative time gain vs. time shows accumulating effect of speed loss, with asymptote towards a total time loss (negative time gain). (**c**) Total time gain vs. up-step height *b* for various nominal walking speeds, with greater bumps causing greater loss of time, and smaller effect for faster nominal speeds. Also shown are effects of late push-off (unfilled symbols). Up-step height *b* = 0.025*L*, nominal mid-stance velocity *V* = 0.44*g*^−0.5^*L*^0.5^. Conditions are equivalent to a human walking at 1.5 m/s, encountering an up-step of 2.5 cm.

The eventual total time loss also increases sharply with up-step height (Fig. 2c). This is because the potential energy of the up-step comes at the expense of (and must not exceed) the model’s kinetic energy, and therefore speed (see velocity 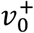; equations 2 and 7). As a result, a faster nominal walking speed is relatively less sensitive to up-step (compare different speeds in Fig. 2c).

The transient response’s decay (Fig. 2a) is well characterized by persistence time. For the nominal gait, the persistence time is about 2.61 steps, equivalent to about 1.39 seconds or 2.06 m for nominal, human-like walking. This is equivalent to a 90% settling time of about 5.7 steps.

### Time may be gained from steady-state walking with a (costly) burst of speed

Given the total time lost from a disturbance, a naïve compensation is simply to regain that time with an isolated time gain, meaning a separate burst of speed from nominal, level walking. The appropriate amount of time *t*_gain_ could be gained either well before or well after nominal speed is disrupted. In either case, the time is to be gained within a burst of *N* fast steps, starting and ending at the nominal steady-state speed *V*, while minimizing total push-off work.

The optimal burst of speed rises smoothly to a single peak, and then decreases back to nominal, with a roughly time-symmetric profile (Fig. 3 top). The corresponding push-off sequence (Fig. 3 top) is neither smooth nor symmetric. The peak push-off work occurs on the initial compensation step, with considerably greater magnitude than nominal, about 1.78 times greater for a sample up-step (*b* = 0.075*L*; *N* = 9). The succeeding push-offs—save the final one—decrease rapidly and then level off above normal, as time is recouped. The final compensatory push-off is actually below nominal, to enable speed-matching for the final conditions. This asymmetric push-off profile nonetheless contributes to a rather smooth accumulation of time gain, resembling a ramp function with rounded corners (Fig. 3 top).

**Figure 3.**
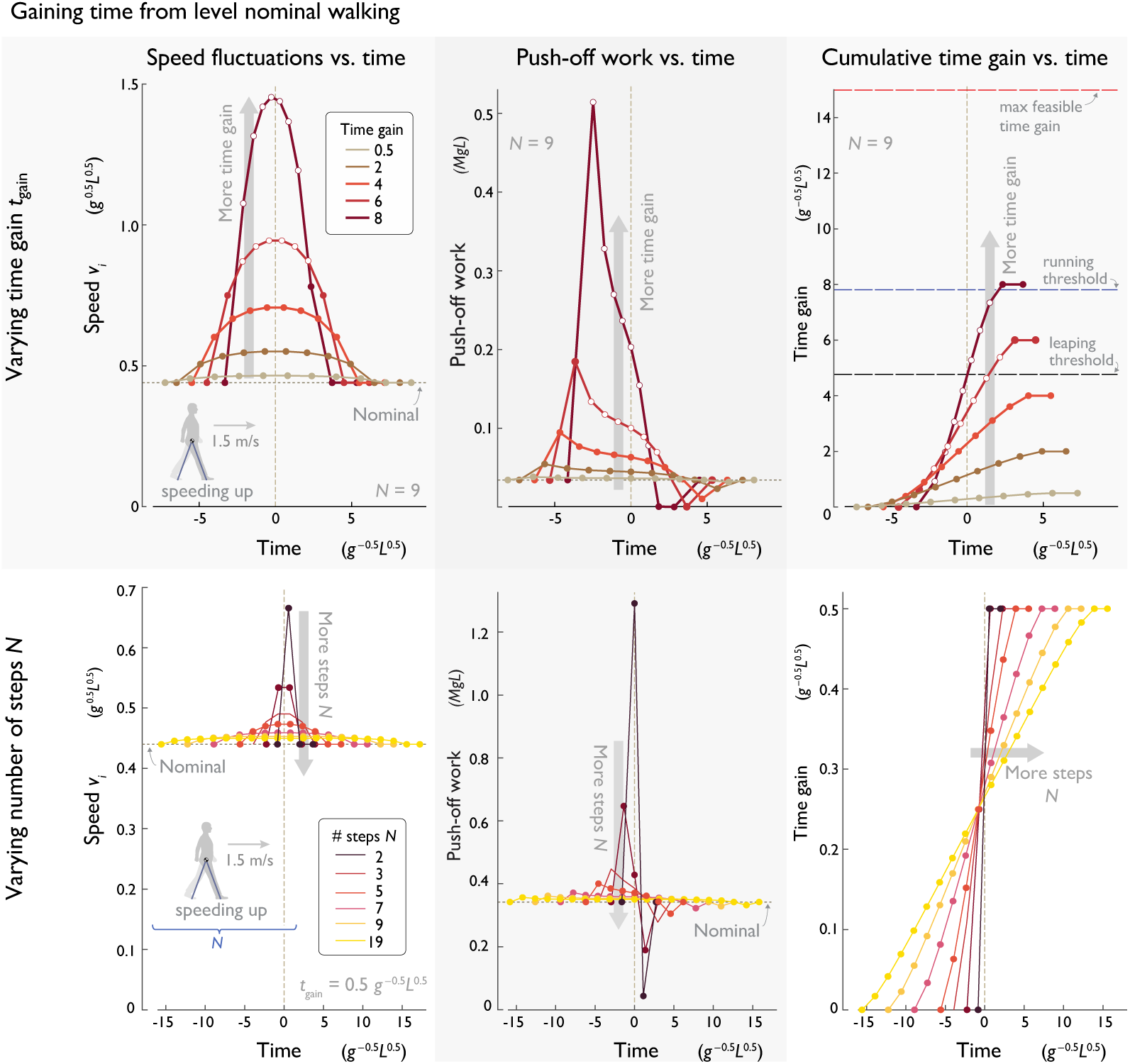
Optimal strategies to gain time during level walking, separate from any disturbance, for (top row:) varying amounts of time *t*_gain_, and (bottom row:) varying number of steps *N*. Each row, from left to right: Speed fluctuations vs. time, push-off work vs. time, and cumulative time gain vs. time, relative to nominal (each step denoted by filled symbol). Parameter values are *N* = 9 steps for top row, *t*_gain_ = 0.5 for bottom row. Filled dots indicate steps. Unfilled dots denote large push-offs that entail momentary aerial phases or even running for several steps; maximum time gain thresholds to avoid such cases are shown in (top most right).

These effects are amplified as more time is gained (Fig. 3 top). The greater the *t*_gain_ (at fixed *N*), the greater both the peak speed and peak push-off. It is also easier to take more compensation steps *N*, which calls for a shallower peak in walking speed, a lower peak push-off, and a more gradual gain, with less time gained per step (Fig. 3 bottom). Conversely, gaining time over very few steps entails a sharp peak in both speed and push-off (Fig. 3 bottom). In terms of both overall and peak amount push-off work, there is great advantage to spreading a time gain over as many steps as possible (Fig. 3 bottom).

Another issue with gaining time is the limitations of a walking gait. A sufficiently large push-off may cause the model to lose ground contact (or leap) into a momentary aerial phase prior to collision (see equation 2), particularly for the first push-offs of the compensation (Fig. 3 top). This might resemble a straight-legged run for a few steps. If such leaping were disallowed, the maximum time that could be gained would be equivalent to about 2.86 nominal step times (*T*). In addition, large enough time gains can call for running, when speed is too high for pendulum-like walking^3^, *v*_*i*_ ≥ 1 *g*^0.5^*L*^0.5^. We did not explicitly model running here, and instead noted which steps or time gains would exceed the thresholds for maintaining a pure walking gait.

The primary issue is the number of steps over which time is to be gained. For fewer steps or more time gain, there are substantial penalties with regard to peaks in walking speed, peak and overall work, and the potential need for aerial phases. For relatively large *N*, the speed and push-off profiles become increasingly constant (Fig. 3 bottom), so that the primary cost is dominated by the work for steady walking at an elevated speed. For large *N*, the extra work needed to gain time is largely inversely proportional to *N*. If time needs to be gained, it should ideally be distributed across as many steps as possible.

### Optimal strategy for an up-step: Speed up, lose speed, speed up

Optimization reveals that the ideal compensation strategy should actually envelop the up-step. A near-optimum may be achieved with a relatively small *N*, with little advantage to taking more steps. Moreover, these *N* steps should optimally be centered about the up-step, with *m* = 0. The best strategy is not to gain time separately from the disturbance, but to modulate push-offs in anticipation of it, and to continue doing so after encountering it. Here we describe the optimal strategies in terms of speed or momentum fluctuations, push-off modulation, and cumulative time gain.

The key to reducing overall work is to ramp up speed (momentum) in a few steps immediately preceding the disturbance (Fig. 4 top), pushing off hardest when stepping onto it. This anticipates a sharp loss in speed atop the up-step, followed by a gradual return to nominal speed, with approximately two-fold symmetric profile: symmetric in time about the up-step, and in speed about nominal speed profile. The associated push-offs increase to a high peak centered on the up-step itself (*u*_0_), and otherwise remain close to nominal for all but a few steps surrounding that step. The push-off profile is also approximately symmetric in time about the up-step, with the exception of the *N*th push-off, which serves a special purpose of matching the final boundary conditions. In terms of cumulative time, the strategy reserves the few final steps to gain time in anticipation of the time lost on and after the up-step. We found little difference to using linearized vs. nonlinear dynamics (solid vs. dashed lines, respectively in Fig. 4 top), and therefore use linearized dynamics for following analyses.

**Figure 4.**
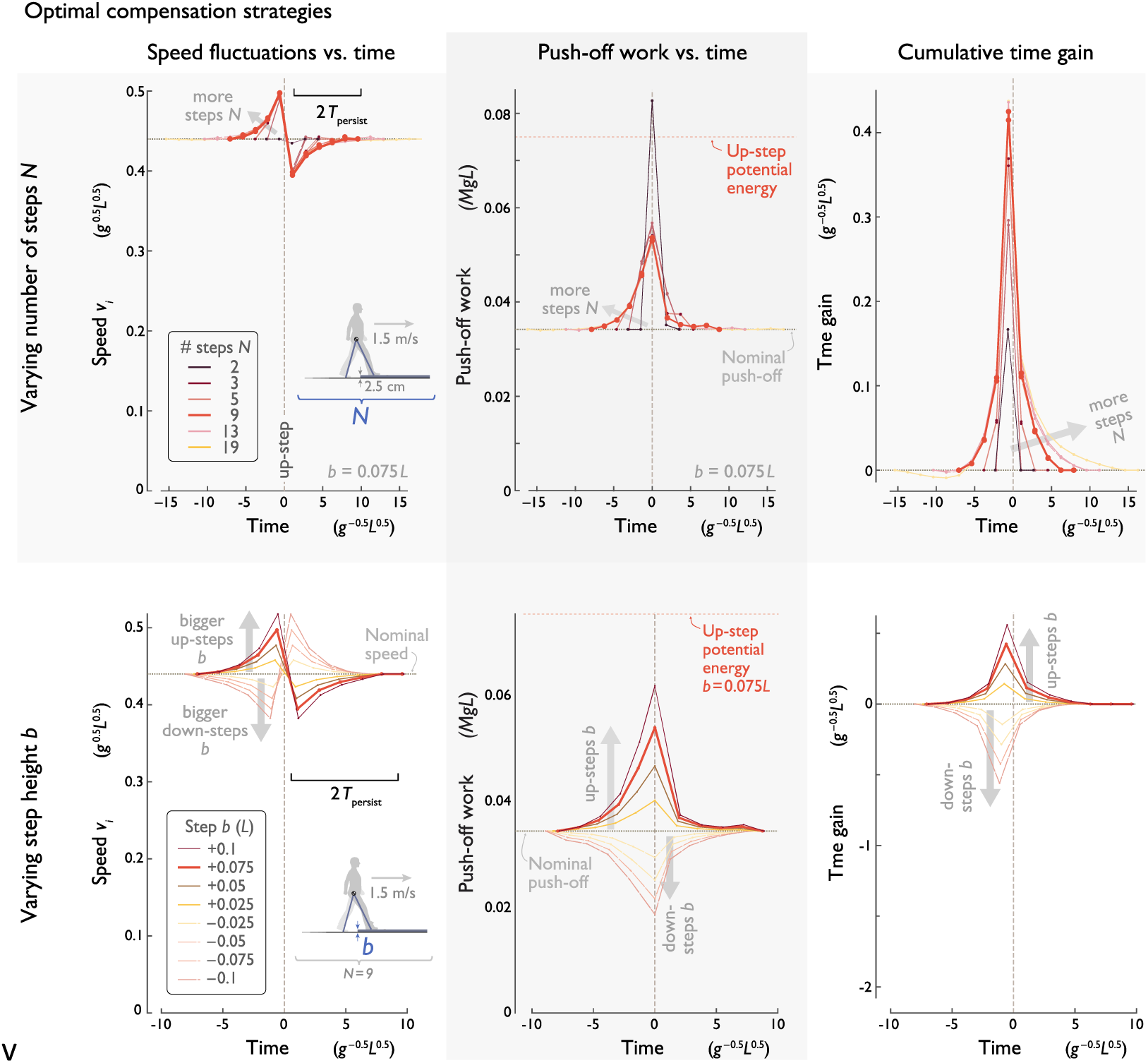
Optimal strategies to compensate for an up-step, for (top row) varying number of steps *N*, and (bottom row) varying step height *b*. Each row, from left to right: Speed *v*_*i*_ vs. time, push-off work vs. time, and cumulative time gain vs. time. In top row, step height is fixed (*b* = 0.075*L*), and optimal solutions shown are for linearized model, superimposed on nonlinear model solutions (dashed lines). In bottom row, both up- and down-step (positive and negative *b*, respectively) solutions are shown for linearized model; *N* = 9. Optimal strategies are all centered about the up-step (*m* = 0).

The compensations increase in magnitude with fewer compensatory steps (Fig. 4 top) or larger up-step heights (Fig. 4 bottom). The extreme case is the minimum possible steps, *N* = 2, which generally call for a large speed-up and a large peak push-off. Not surprisingly, the amplitudes of both speed and push-off profiles increase with bump height. Moreover, the profiles scale approximately linearly for small *b*. But regardless of the number of steps and the height to be gained, the optimal compensation largely occurs within ±2*T*_persist_ of the up-step.

The optimal strategy takes unique advantage of the dynamics of the up-step. There is a unique reduction of kinetic energy exchanged for the up-step’s potential energy, which accounts for a substantial loss of speed during the single upward step. That transient loss is compensated for by speeding up in two bouts, before and after the up-step. Speeding up is economical because the larger push-offs result in reduced collisions. It is optimal to mostly concentrate loss of speed into a single step, and to distribute the accelerations across many steps. This is illustrated by the case of many steps (e.g. see Fig. 4 top, *N* = 19), where there is slight advantage to first slowing down slightly, to extend the speed-up prior to the up-step.

We additionally observe that the optimal compensation for a down-step (Fig. 4 bottom), largely mirrors that for an up-step. The actions largely scale with step height, so that negative heights (down-steps, *b* < 0) entail slowing down rather than speeding up ahead of the disturbance and reducing push-offs instead of increasing. Similarly, the cumulative time change is also nearly opposite to an up-step, except with time being deliberately lost instead of gained, in anticipation of the kinetic energy gained from stepping down. This scaling effect occurs about the nominal speed, push-off, or time gain, and is not quite linear, owing to the nonlinear dynamics (equations 2 & 9). Nevertheless, scaling of compensation with step height *b* serves to roughly describe the optimal solution for both up- and down-steps. Down-step strategies will be examined further in another section below.

### Earlier or later compensations are more costly

The optimality of centering the compensation about the up-step is illustrated by the costs of doing otherwise (Fig. 5). Shifting the compensation earlier or later (*m* < 0 or *m* > 0, respectively) and optimizing for each case, reveals increasing costs for total push-off work (Fig. 5a) and peak push-off work (Fig. 5b).

**Figure 5.**
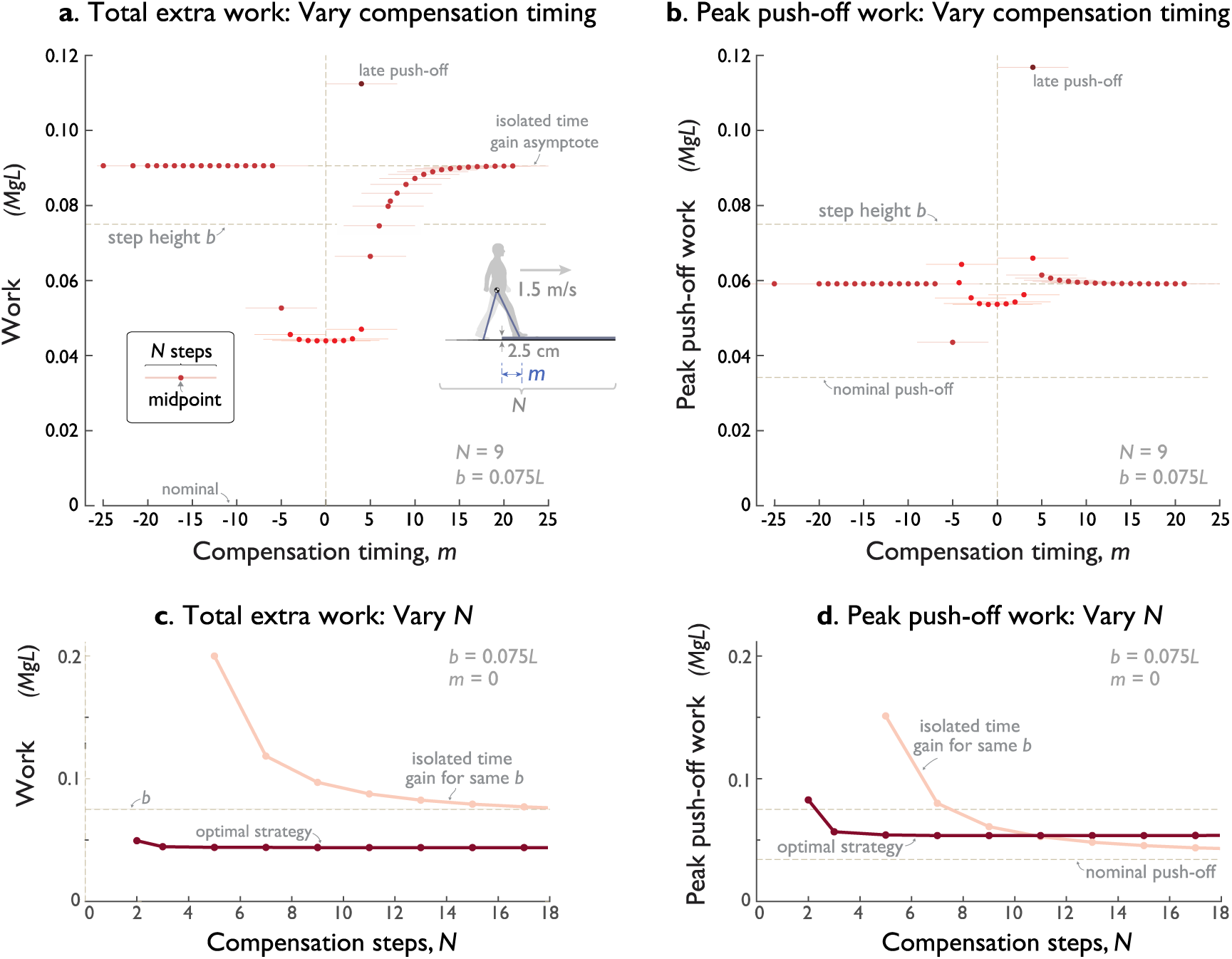
Costs for optimal compensation strategies, as a function of compensation timing (shifting by *m* relative to up-step, see Fig. 1) and the number of compensation steps (*N*). (**a**) Total extra work vs. *m*, for *N* compensatory steps (*N* = 9, *b* = 0.075*L*). Total extra work is defined as the work in excess of *N* nominal steps. Filled symbols indicate cost for each *m*, lighter horizontal lines indicate compensation range of *N* steps. (**b**) Peak push-off work vs. *m*. (**c**) Total extra work vs. *N*, and (**d**) Peak push-off work vs. *N* for optimal compensation centered about up-step (*m* = 0). Also shown is the cost of an isolated time gain from nominal walking (Fig. 3), to regain time lost from an equivalent up-step (*b* = 0.075*L, t*_gain_ = 2.48 *g*^−0.5^*L*^0.5^) with no compensation. In (**a**) and (**b**), minima for both total and peak work occur at *m* = 0, where compensatory steps are distributed equally before and after the disturbance. Both costs also decrease for greater *N*. Also shown in (**a**) is the optimal strategy for recovering from an unanticipated up-step with a late push-off on the up-step.

There is still more cost if the compensation occurs entirely before or after the disturbance. Compensating early with no overlap 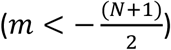 is essentially equivalent to a completely separate time gain (Fig. 3), with an equivalent higher cost. Compensating later 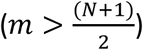 also tends toward the same higher cost, albeit with a longer transition. This is due to the transient response after the disturbance, in which the total time loss accumulates over several persistence times *τ*. Beginning the compensation during the transient therefore means that less time is to be regained, and so the overall work cost only gradually increases towards an asymptote (Fig. 5a), namely the cost for gaining time separately. The peak push-off also tends toward its corresponding asymptote (Fig. 5b), albeit with elevated costs in some cases. These transition cases reinforce that the optimal compensation should envelop up-step symmetrically.

A large number of steps is not necessary to obtain near-optimal total work or peak push-off work. Although both costs (Fig. 5c and d) decrease with greater *N*, the returns diminish rapidly. For example, taking the minimum of two compensatory steps only takes about 12% more extra work than for nine (*N* = 9; *b* = 0.075*L*). Beyond about two persistence times (2*T*_persist_), the costs are nearly indistinguishable. There is little to be gained from looking more than a few steps ahead.

The optimal strategy has considerable advantages over an isolated time gain, separate from the up-step. For the sample comparison (Fig. 5), the optimum overall work is roughly half that for a completely separate time gain (compare minimum with asymptote in Fig. 5a). There is also a high cost penalty for failing to anticipate the up-step at all, and starting the compensation after a late push-off^17^. A similar but smaller advantage is observed in peak push-off. However, there is a special case of a compensation that ends just before the up-step (*m* = −(*N* + 1)/2 in Fig. 5b), where peak push-off is reduced, albeit at a slight cost in total work.

It is interesting to note that the optimal strategy is generally less costly than the potential energy of the up-step itself. One might expect a cost equal to the work for level nominal walking, plus the work needed to lift the body up the step. But here (Fig. 5), the overall added cost is about 58.5% of the potential energy gain, and the peak push-off is about 71.5%. As stated earlier, the optimal strategy actually gains advantage from the up-step.

### Optimal strategies are self-similar despite parameter variations

The optimal strategies have similar speed profiles even for different parameter values for average speed and step length (Fig. 6). Taking greater (fixed) step lengths while keeping average walking speed fixed, the optimal compensations have similar shape but in fewer steps and less amplitude (Fig. 6a). The speed profiles are also similar to each other when increasing speed while keeping step length fixed (Fig. 6b), although the amplitudes decrease. Thus, for a wide range of parameter values, the compensation strategy always calls for speeding up ahead of an up-step, losing speed atop the up-step, and then recovering speed for several steps after it.

**Figure 6.**
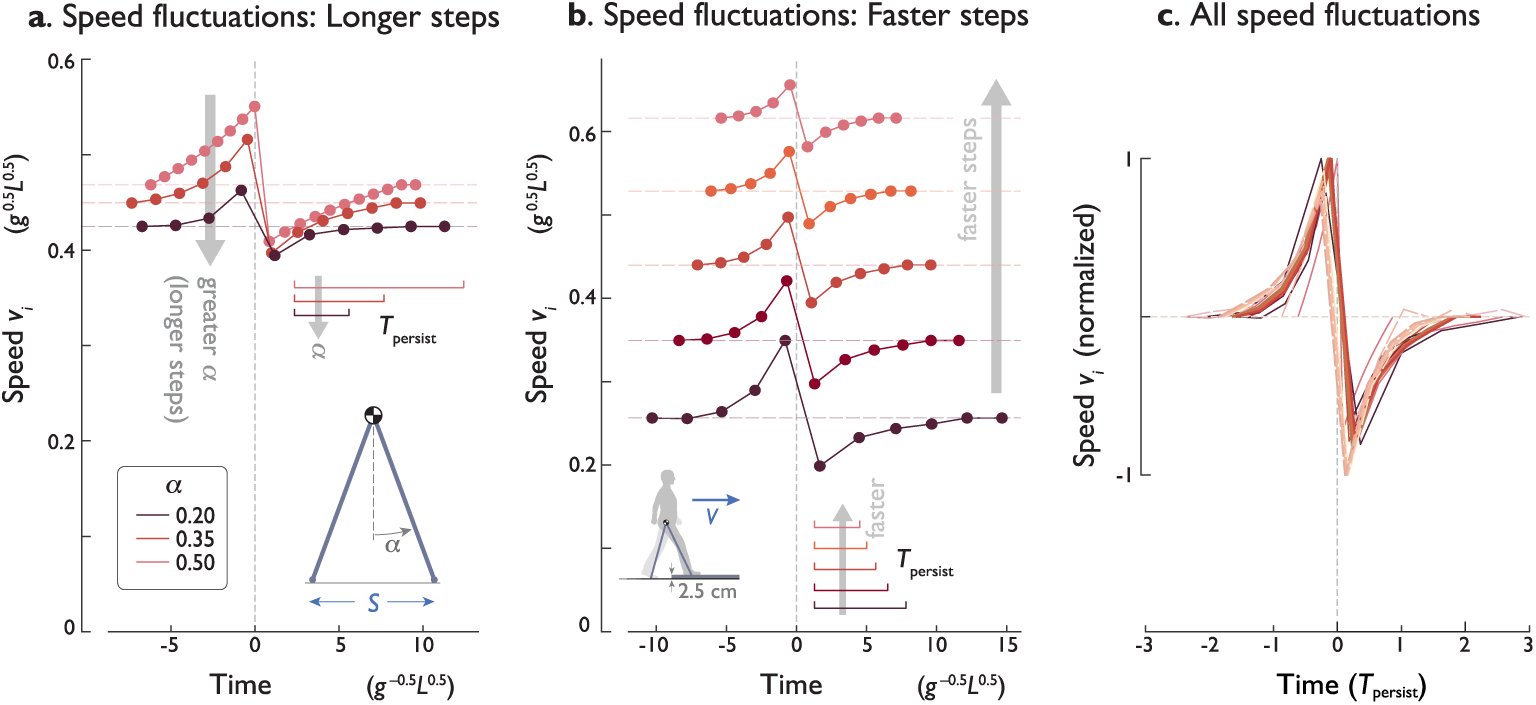
Effect of parameter variations on optimal strategies. (**a**) Optimal speed fluctuations for different values of step length parameter (via leg angle *α*; constant average speed). (**b**) Optimal speed fluctuations for different average speeds (via step frequency; constant leg angle *α*). In both (**a**) and (**b**), *b* = 0.075*L* and *N* = 9. (**c**) Speed fluctuations for all parameter variations considered thus far, scaled by persistence time. Superimposed are variations in number of steps *N* (Fig. 4 top), step height *b* (Fig. 4 bottom), step length, and speed. All amplitudes are plotted relative to respective nominal speed, and have been normalized to unity positive peak. Speed fluctuations for negative step heights are shown negated to facilitate comparison.

All of these effects may be summarized in terms of persistence time. Whether varying speed or step length, the optimal compensation maintains approximately the same profile (albeit varying amplitude) and takes place within about ±2*T*_persist_. In fact, including all parameter variations considered thus far (e.g., number of steps, step height), the speed fluctuation profiles are nearly the same when normalized by persistence time (Fig. 6c). Persistence time is helpful for summarizing the combined effects of a discrete, once-per-step collision, and the continuous-time dynamics of the pendulum-like stance leg. It also plays a large role in the optimal compensation, whether stepping up or down, or walking at a range of speeds and step lengths.

### Up- and down-step strategies are related due to time-symmetry

The optimal strategy to compensate for a down-step is a simple time-reversal of the up-step strategy (Fig. 4 bottom). The speed profile for the down-step (*b* < 0) is the same as for the up-step except with time running backwards. In fact, the push-off profile for a down-step is the same as the time-reversed up-step profile, except using the up-step collisions as the down-step push-offs. Similarly, the cumulative time gain is reversed in time, where down-step time loss is equivalent to up-step time gain.

Time-reversal symmetry is explained by two basic principles. First, the walking dynamics have time-symmetry. The push-off and collision impulses act the same and differ only in which leg they apply to (trailing and leading leg, respectively), and in the order of execution. As a result, the same dynamics may be played forward or backward in time, and the model walks in one direction or the other (to right or left of Fig. 1). For example, the push-off profile for a down-step (e.g., *b* = −0.05*L* in Fig. 4 bottom) may be flipped about the (vertical) work axis (at *t* = 0) to yield the collision profile for an equivalent up-step (*b* = 0.05*L*).

The second principle is that minimizing push-offs and minimizing collision magnitudes yield the same solution. This is because the boundary conditions enforced here call for nominal level walking at the same speed, and thus no difference in kinetic energy between start and end. The only energy difference is in the potential energy of the step (*Mgb*). The only energy sources and sinks are the push-offs and collisions, and so the sum of all push-offs and collisions must equal the potential energy, which is fixed for a given optimization problem. In addition, push-offs can only perform positive work, and collisions negative work. Any solution that minimizes total push-offs must therefore also minimize the total collision magnitudes.

Next is to apply time-symmetric dynamics to the optimization. The optimal strategy to traverse an up-step with minimal push-off work (e.g., Fig. 4 top) may be played in reverse, to yield the optimal strategy to traverse a down-step with minimal collision work magnitude. But minimizing collision magnitude is the same as minimizing push-off, and so the same strategy, whether played forward or backward in time, minimizes the respective push-offs for either direction.

## Discussion

We explored the optimal strategy for anticipating and compensating for a single up- (or down-) step on otherwise flat terrain. The uncompensated effect of the disturbance was characterized in terms of a transient response with a loss of time and forward momentum, followed by an asymptotic return to nominal walking. For planned tasks, it may be desirable to regain the lost time, which was examined in two cases: one where time is regained by speeding up level walking, and the other, optimal case, where the compensatory steps are distributed both before and after the step. The analysis yields several conclusions regarding optimal compensations.

One conclusion is that there is a single, consistent pattern to optimally compensating for an up-step. That pattern consists of speeding up prior to the up-step, losing speed (and time) ascending that step, and then speeding up again after it (Fig. 4). With appropriate scaling, that basic pattern applies regardless of the number of compensation steps, the up-step height, the walking speed, or the step length (see Fig. 6). For example, the pattern’s timing scales largely in accordance with the persistence time, which is determined by the speed and step length (see Methods, equation 10). The pattern’s amplitude also scales, for different heights (including down-steps), speeds, or step lengths, but relatively little with the number of compensation steps. In fact, although there is always an advantage to taking more steps, there is rather little advantage beyond *N* = 3 steps. A general rule of thumb is that the optimal compensation should largely occur within ±2 persistence times of the up- (or down-) step.

This strategy is also clearly superior to gaining time separately during level walking. For example, the optimal compensation in five steps requires 51.3% less total extra work and 8.5% less peak work (*N* = 5, *b* = 0.075*L*; see Fig. 5), while also having less sensitivity to the number of compensation steps. This is because a separate time gain is largely a matter of walking faster, which is costly on level ground. But the transient loss of speed and time from the up-step allows opportunity to reduce collision losses over two bouts of acceleration. Less energy is lost to collisions when either accelerating or stepping upward, and the optimal compensation takes full advantage of such opportunities.

These results help to reinforce the benefits of looking ahead. Vision (or other imaging) is certainly critical for spatial path planning, for example to determine feasible paths to avoid obstacles. But the dynamics of legged locomotion are such that planning can also be performed dynamically, in terms of momentum trajectories. Knowledge of upcoming terrain variations can allow not only for economy, but also for surmounting obstacles that might otherwise exceed actuator limitations. An up-step can thus be surmounted with less loss of time, less overall work, and reduced peak push-off compared to the up-step’s gravitational potential energy.

This has potential implications for control of legged robots. Perhaps the most straightforward control for robots, and also the most important for stability, is state feedback control. Such control might automatically regulate walking speed toward a nominal reference value, but would not normally be expected to make up for lost time, unless time were included as an explicit goal, and perhaps included as an explicit state (e.g., integrated error in the feedback controller). Even then, responding after a disturbance (see separate time gain, Fig. 3; or later timing, Fig. 5) cannot be as economical as the optimum (*m* = 0, Fig. 5). Although the true optimum appears to require infinite look-ahead, near-optimal performance is possible with only about two persistence times (or distances) of foresight. Although such compensation could be stored as a scalable trajectory (e.g., Fig. 6c), an alternative would simply be a model predictive control^18^, where an optimal plan is continually regenerated with a receding horizon. That horizon need only be about two persistence times ahead, and might be compatible with other optimization approaches already employed for some legged robots^19–22^.

Our analysis may suggest how humans negotiate small obstacles. Humans rely on vision to identify curbs or height discrepancies in a sidewalk, and the resulting compensations may do more than simply prevent tripping. It is possible that humans speed up prior to a stepping up a curb, and perhaps slow down prior to stepping down a curb^23^. They appear to look several steps ahead^24^ and make quick, dynamically sensible decisions. Moreover, they appear to control not only speed but also foot placement, while taking into account actuator limitations. It is unknown how humans actually plan their momentum or whether they control it optimally. The optimal strategies considered here might serve as a reference for comparison to humans.

The compensation strategies examined here are focused on pendulum-like walking. Our step-to-step transition analysis treats the leg as relatively straight when it contacts ground. This may apply well to relatively small up-steps, but not to larger ones. Humans normally flex the swing knee at mid-stance, in part to gain ground clearance, and then extend again prior to ground contact^25^. But swing knee extension does not seem to apply for larger inclines or typical stairs, where there may be less forward momentum. Compensation strategies might be different for larger up-steps, and especially for less pendulum-like gaits.

The present analysis has a number of additional limitations. Our model has fixed step length and width, and performs impulsive step-to-step transitions, whereas humans actually modulate step distances, and perform step-to-step transitions over finite time. The actual costs of varying speed^26^ might therefore differ from that considered here. The model is also confined to dynamical planning of push-offs and speed, assuming that the spatial path is already determined. An improved model could allow for three-dimensional stepping^27,28^ with multiple joints that allow for leg compliance^29^ and finite-duration step-to-step transitions, and might combine spatial and dynamical planning into a single optimization problem.

Despite these limitations, there may be some general conclusions that may yet apply to more complex models. Even with additional degrees of freedom, walking is often dominated by forward momentum, and the step-to-step transition may dissipate significant energy, which must then be restored with active work. It may therefore be advantageous to counter a step height disturbance by gaining momentum several steps beforehand and continuing the compensation past the disturbance for several steps thereafter. Furthermore, the appropriate compensation range may be described by a measure such as persistence time. Without planning, compensations for uneven terrain may come with a relatively high cost in overall work, peak actuator demands, and/or number of compensation steps. There may be considerable advantage to applying optimization to the dynamics of legged locomotion, particularly on uneven terrain.

## Methods

### Model of walking

The walking model has rigid, pendulum-like legs whose ground collisions dominate the energetics (Fig. 1a). It has a point-mass pelvis of mass *M*, atop massless legs taking fixed-length steps (Fig. 1b). The stance leg behaves like an underactuated, inverted pendulum during single support, punctuated by the “step-to-step transition” to redirect the center-of-mass (COM) velocity between steps^14^. This is accomplished with an active push-off along the trailing leg, followed immediately by an inelastic heel-strike collision along the leading leg, both modeled as ideal impulses in immediate succession (Fig. 1c). A full walking step therefore starts with a push-off then a collision and followed by pendular stance phase that ends just before the next leg contacts ground. Middle stance (mid-stance) is defined as the instant that the stance leg is upright within the stance phase. When the terrain disturbance is encountered (Fig. 1d), we consider cases where push-off occurs pre-emptively or late, relative to collision. In the optimal case (Fig. 1e), terrain variations are sensed in advance, and pre-emptive push-offs are to be planned. We previously examined a similar model^17^, and found that failure to anticipate and perform push-off pre-emptively, for example on uneven terrain, results in considerably poorer economy.

The model dynamics are briefly summarized as follows (detailed previously^17^): Each step has index *i* with the disturbance located at *i* = 0 (Fig. 1d). Negative *i* therefore refer to the preparatory steps beforehand, and positive to recovery steps thereafter. Each step begins just before the step-to-step transition, with COM velocity 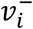 directed forward and downward as dictated by the preceding step’s stance phase. The ideal step-to-step transition starts with pre-emptive push-off work *u*_*i*_ (in units of mass-normalized work) performed impulsively along the trailing leg to redirect the COM velocity. Applying impulse-momentum (and normalizing by body mass) yields an intermediate velocity of magnitude 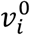,

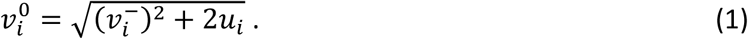

This is followed immediately by the heel-strike collision along the leading leg, to yield post-collision velocity 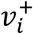. Again applying impulse-momentum,

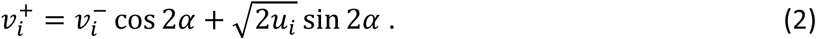

The single stance phase follows the step-to-step transition, and is modeled as an underactuated, simple inverted pendulum. As a discrete measure of overall forward momentum, we use the mid-stance velocity *v*_*i*_ (no superscript; see Fig. 1e), sampled when the leg is vertical and the COM velocity is purely forward.

We treat steady, level walking as the nominal condition (Fig. 1c). Each begins and ends at the same speed 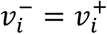. The nominal push-off work *u*_*i*_ offsets the collision work^14^, so that

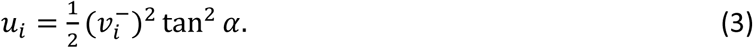

The up- or down-step disturbs steady walking (Fig. 1d). Its height *b* (positive for up-steps, negative for down-steps) causes the preceding step (*i* = −1) to end with different leg angles from nominal. For simplicity, we will typically refer to up-steps alone, and treat down-steps as a straightforward generalization there-of, unless otherwise discussed. For a given height *b* and step length *S*, we define the angular disturbance as *δ*_*i*_,

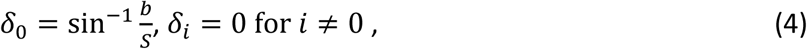

where the angle is zero for all non-disturbance steps.

We expect that it is helpful to detect the up-step ahead of time, to allow push-off to occur pre-emptively, that is, before the collision. Pre-emptive push-off is more economical both on the level^14^ and on the up-step^17^, because it reduces the subsequent collision loss. If the up-step is completely unanticipated, the push-off might occur late. If so, the leading leg collision first redirects COM velocity along a new pendular arc. We refer to the work of a late push-off as 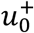 (with the plus sign ‘+’ indicating the post-collision timing), performed impulsively to increase speed along that new arc:

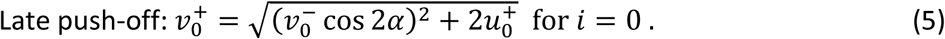

We treat the late push-off as of nominal amount, as if the nominal energy were stored elastically as observed in humans^30,31^, but released late. The next step will then have substantially reduced speed 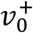 and slower step time *τ*_0_.

The inverted pendulum stance phase follows the step-to-step transition. The step time *τ*_*i*_ is defined as the time for the stance leg angle *θ* to move from initial to final angles (*α* + *δ*_*i*_ and −*α* + *δ*_*i*+1_, respectively). Using the linearized dynamics,

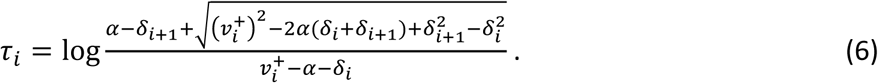

Solving the equation of motion with the step time, the velocity at end of stance 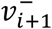, or equivalently the beginning of the next step-to-step transition can be found as:

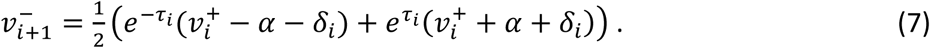

We chose nominal parameters to correspond to typical human walking. A person with leg length *L* of 1 m may typically walk at 1.5 m/s, with step length of 0.79 m and step time of 0.53 s (from anecdotal observations). Using dynamic similarity, parameters and results may be expressed in terms of body mass *M*, gravitational acceleration *g*, and *L* as base units. The corresponding model parameters are angle *α* = 0.41, push-off *U* = 0.0342 *MgL*, step time *T* = 1.66 *g*^−0.5^*L*^0.5^, and pre-collision speed *V*\* = 0.601 *g*^0.5^*L*^0.5^, where capital letters indicate nominal values for *u*_*i*_, *τ*_*i*_, and 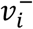, respectively. We also refer to a nominal speed *V* = 0.44 *g*^0.5^*L*^0.5^ for mid-stance speed *v*_*i*_. We considered a range of up-step heights, for example *b* = 0.075*L*, equivalent to about 7.5 cm for a human.

Although most of this study relied on the linearized dynamics of equations (6-7), we also performed a subset of simulations with fully nonlinear dynamics to test the accuracy of this approximation (Fig. 4).

### Optimal Control Formulation

We used optimization to determine the push-off sequence that modulates momentum to most economically negotiate a single disturbance. The objective was to minimize the total positive work of *N* consecutive push-offs of amount *u*_*i*_, subject to constraints to depart from and then regain nominal walking with no loss of time or speed compared to steady, level gait. The timing of this compensation was found to affect overall economy and was therefore explored by shifting the *N* modulated steps earlier or later relative to the disturbance. The optimal strategies were explored with parameter variations in compensatory steps *N*, step height *b*, nominal speed *V*, and compensation timing *m*.

The *N* compensatory steps were nominally centered about the up-step. It is convenient to consider odd *N*, so that the single up-step at *i* = 0 is surrounded by an equal amount of anticipatory steps before, and recovery steps after. The parameter *m* (Fig. 1e) shifts these steps later (*m* > 0) or earlier (*m* < 0). Again for odd *N*, the compensation steps range from 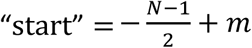 to 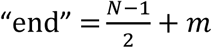. The speed constraint states that walking speed begins at and returns to nominal, level walking. The time constraint requires that the total time for stepping onto and recovering from the uneven terrain, be equal to the nominal time to walk the same number of level steps.

The optimal policy *π* (sequence of push-offs *u*_start_, …, *u*_end_, all constrained to be non-negative) is:

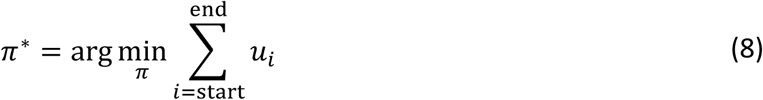

Subject to:

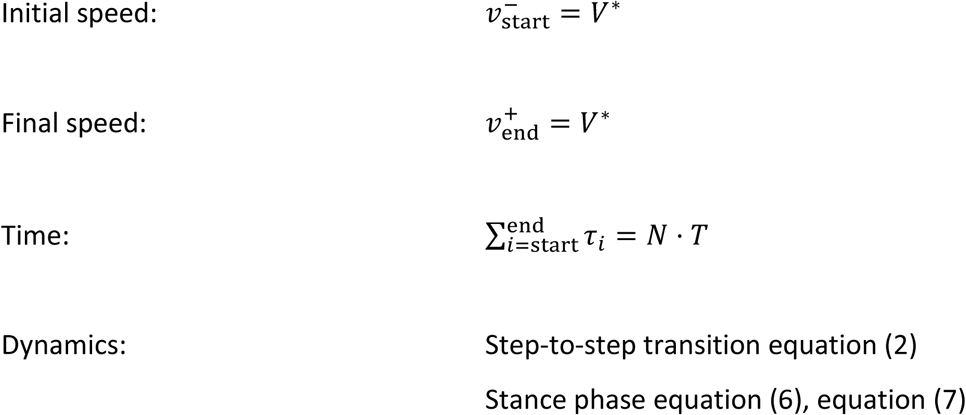

We next define a persistence time for the hybrid dynamics of discrete step-to-step transitions and continuous-time pendulum dynamics. On level ground and with unmodulated push-offs, walking speed converges exponentially toward nominal with each discrete step. The step-to-step eigenvalue cos 2*α* (about 0.68 for nominal gait; equation 2) determines this convergence. This may be combined with the continuous-time step period *T* to yield an overall decay with time constant *T*_persist_:

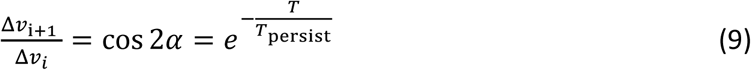

where Δ*v*_*i*_ refers to the deviation from nominal mid-stance speed at step *i*. Solving for persistence time,

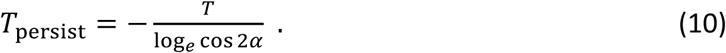

An analogous persistence distance may be found by dividing *T*_persist_ by nominal speed. Persistence time (or distance) may be interpreted as the time (or distance) required for a disturbance response to decay to *e*^−1^ ≈ 37% of its peak amplitude. It helps to place an expectation on optimal recovery from a disturbance. With active control of push-off, it is of course possible to change the disturbance response substantially. But when optimizing for work or energy, we do not expect the response to be far faster than uncompensated, because that would entail a high effort cost. Nor do we expect the response to be slower than the uncompensated case, which is already stable. We therefore expect that optimal compensation responses should be somewhat faster than uncompensated and should therefore decay within no more than a few persistence times.

## Acknowledgements

This work is supported by NSF DGE 0718128, the ONR ETOWL program, NIH AG030815, the Dr. Benno Nigg Research Chair (University of Calgary), and NSERC (Natural Sciences and Engineering Research Council of Canada) Discovery program and Canada Research Chair (Tier 1) program.

## Author Contributions

O.D. performed the simulations and generated the figures. O.D. and A.D.K. wrote the manuscript. A.D.K. also revised the manuscript and the figures. H.T. reviewed the manuscript.

